# Peptidome profiling for the immunological stratification in sepsis: a proof of concept study

**DOI:** 10.1101/2020.12.29.424724

**Authors:** Martín Ledesma, María Florencia Todero, Lautaro Maceira, Monica Prieto, Carlos Vay, Marcelo Galas, Beatriz López, Noemí Yokobori, Bárbara Rearte

## Abstract

Sepsis has been called “the graveyard of pharmaceutical companies” due to the numerous failed clinical trials. The lack of tools to monitor the immunological status in sepsis constrains the development of immunomodulatory therapies. Here, we evaluated a test based on whole plasma peptidome acquired in a MALDI-TOF-mass spectrometer used for bacterial biotyping and machine-learning algorithms to discriminate the different immunological phases of sepsis. In this proof of concept study, two discrete lipopolysaccharide-(LPS) induced murine models emulating the pro- and anti-inflammatory phases that occur during sepsis were evaluated. The LPS group was inoculated with a single high dose of LPS, recalling the proinflammatory phase, and the IS groups was subjected to increasing doses of LPS to induce the anti-inflammatory/immunosuppression phase. Unstimulated mice served as controls. Both experimental groups showed the hallmarks of pro- and anti-inflammatory phases respectively; the LPS group showed leukopenia and higher levels of cytokines and tissue damage markers, and the IS group showed neutrophilia, lymphopenia and significantly lower antibody titers upon immunization. Principal component analysis of the plasma peptidomes formed three discrete clusters that mostly coincided with the experimental groups. In addition, machine-learning algorithms discriminated the different experimental groups with a sensitivity and specificity of up to 95.7% and 90.9% respectively. Our data reveal the potential of plasma peptidome analysis by MALDI-TOF-mass spectrometry as a simple, speedy and readily transferrable method for sepsis patient stratification that would contribute to therapeutic decision-making based on their immunological status.

## 1 Introduction

Sepsis constitutes one the major causes of death in intensive care units (Vincent et al., 2009; Novosad et al., 2016) with an incidence of 31.5 million cases and 7-9 million deaths annually (Fleischmann et al., 2016). The impact of sepsis in developing regions such as Latin America is elusive due to the limited number of prevalence studies, but the high incidence rate of infectious diseases and the inequities in the access to health care indicates that sepsis is an alarming health problem in the region (Machado et al., 2017; Azevedo et al., 2018; Estenssoro et al., 2018; Estenssoro et al., 2019).

The redefinition by the Sepsis-3 consensus highlights the need to integrate the information on the etiological agent and the host response status and strongly emphasizes that the dysregulated host response as a key component of sepsis (Singer et al., 2016).

From the immunological point of view, sepsis is characterized by the predominance of an overwhelming pro-inflammatory response during the early stages with a concomitant anti-inflammatory and immunosuppressive response that becomes predominant in the later stages (van der Poll et al., 2017; Venet et al., 2018; Rubio et al., 2019). Sepsis has earned the epithet of “the graveyard of pharmaceutical companies” due to the numerous failed clinical trials and this could be related to an inappropriate timing for the application of immunomodulatory interventions (Rubio et al., 2019). The lack of tools to monitor the host immune status constrains the evaluation of novel immunomodulatory therapeutic approaches. Despite the efforts put in biomarker search in the past decades, only a handful of molecules, such as procalcitonin, proved to be useful in the clinical setting under specific conditions, and the fact that efficient biomarkers not necessarily correspond to key mediators adds further complexity (Parlato and Cavaillon, 2015). Moreover, none of them determines the quickly changing immunological status of the host (Pierrakos and Vincent, 2010; van der Poll et al., 2017; Parlato et al., 2018; van Engelen et al., 2018b; Venet et al., 2018; Al Jalbout et al., 2019; Gunsolus et al., 2019; Rubio et al., 2019; Schenz et al., 2019).

With the advent of artificial intelligence algorithms including machine learning (ML) (López Fernández et al., 2016), top-down analysis of unlabeled biological samples is gaining ground. This is the case of matrix-assisted light desorption ionization-time of flight-mass spectrometry (MALDI-TOF-MS), a powerful tool that is currently spreading in the clinical setting for microbiological biotyping and in biomedical research for the study of biological samples in several diseases including sepsis (Ludwig and Hummon, 2017; Hou et al., 2019).

Taking into account that around 50% of all sepsis cases are caused by Gram-negative bacteria (van Engelen et al., 2018a; Dolin et al., 2019), we (Rearte et al., 2010a; Rearte et al., 2010b; Landoni et al., 2012; Martire-Greco et al., 2014; Rearte et al., 2014; Córdoba-Moreno et al., 2019; Montagna et al., 2020) and other authors (Opal et al., 1999; Genga et al., 2018) demonstrated that both inflammatory and anti-inflammatory processes that occur during sepsis can be emulated by lipopolysaccharides (LPS) in murine models.

Herein, we hypothesized that the pro/anti-inflammatory phases could be discriminated through the analysis of plasma peptidome spectra generated by MALDI-TOF-MS. Thus, in this study, using LPS-induced murine models emulating the different phases, we developed and evaluated predictive models based on ML algorithm that would allow the discrimination of the different immunological phases.

## 2 Materials and methods

### 2.1 Animals

Female and male BALB/c mice (8–12 weeks old) were provided by the IMEX-CONICET-Academia Nacional de Medicina, Buenos Aires, Argentina. Animals were maintained under a 12 h light–dark cycle at 22 ± 2 °C and were fed with standard diet and water *ad libitum*. Animals were bred and housed in accordance with the NIH Guide and Use of Laboratory Animals (National Research Council (U.S.), 2011). Experimental designs were approved by the Committee for the Care and Use of Laboratory Animals (CICUAL) of IMEX-CONICET, Academia Nacional de Medicina de Buenos Aires.

### 2.2 Murine Models

All the inoculation and sample collection schemes are depicted in Supplementary Figure 1.

Briefly, for the inflammatory phase model (LPS group), BALB/c mice were inoculated with a lethal dose 50 (LD50) of LPS of *Escherichia coli* O111: B4 (100 μg/ mouse; Sigma-Aldrich, St Louis, MO, USA) intraperitoneally (i. p.) (Córdoba-Moreno et al., 2019). Plasma samples were obtained at 1.5 h and 6 h after the injection. In the anti-inflammatory/immunosuppression phase model (IS group), BALB/c mice were inoculated daily with LPS for 10 consecutive days. The inoculation scheme consisted of increasing doses starting from 5 μg/mouse i. p. for the first 4 days, followed by 50 μg/mouse i. p. for 3 days, and 100 μg/mouse i. p. for the last 3 days (Rearte et al., 2010b). Plasma samples were obtained 24h after the last LPS dose. A third group of mice inoculated with vehicle (saline solution) served as control (CTL group). The plasma in this group was collected at the same time points indicated in the experimental groups. Peripheral blood was collected by submandibular bleeding method in order to maximize the quality of the sample. Part of the heparinized samples were used for blood cell count and the remaining volume was centrifuged twice (400xg, 10 min at 4°C) and plasma were stored at −20°C until analysis.

### 2.3 Immunological, hematological and biochemical parameters

Peripheral blood leukocytes were counted in a Coulter hematology analyzer (Diatron Abacus Junior Vet, Budapest, Hungary) at 1.5 h and 24 h post LPS for the LPS and the IS groups respectively. Proinflammatory (TNF-α; IL-12p70; IFN-γ; IL-6) and anti-inflammatory cytokines (IL-10, TGF-β) were determined in plasma by ELISA at the indicated time points (OptEIA set; BD Biosciences, San Diego, CA, USA) according to the manufacturer’s instructions. The tissue damage markers creatinine (Cre), alanine aminotransferase (ALT) and aspartate transaminase (AST) were determined in plasma, at 6 and 24 h in the LPS and IS groups respectively, using a kit from BioTecnica (Varginha, Minas Gerais, Brazil) and the MINDRAY BS-200E auto-analyzer according to the manufacturer’s instructions.

For flow cytometry analysis, 24 h after the last LPS dose, a splenocyte suspension was obtained and the cells were immunolabeled as described previously (Rearte et al., 2010b; Rearte et al., 2014). The following cell types were evaluated: CD4 (FITC-anti-CD4, clone RM4-5) and CD8 (PECy5-anti-CD8, clone 53-6.7) T-lymphocytes, B-lymphocytes (FITC-anti-CD45R-B220, clone RA3-6B2), polymorphonuclear neutrophils (FITC-anti-CD11b, clone M1/70 and PE-anti-Ly6G, clone 1A8). The expression levels of PDL-1 (PE-anti-PDL-1, clone MIH5) on the CD11b gate of myeloid lineage cells were also evaluated. Labeled monoclonal antibodies were obtained from Invitrogen and BD Pharmingen™. Cells were acquired in a Becton Dickinson FACScan flow cytometer using Cell Quest software (Becton Dickinson, San Jose, CA, USA).

Primary antibody response to sheep red blood cells (SRBC) was evaluated by immunizing animals 24 h after the last dose of LPS (5×10^8^ SRBC /mouse, 0.1ml i. p.). The anti-SRBC antibody titer was evaluated in serum through an hemagglutination assay 7 days after immunization as described previously (Rearte et al., 2010b; Rearte et al., 2014).

### 2.4 Acquisition of MALI-TOF-MS spectra

Plasma samples were analyzed with the MALDI Biotyper System (Bruker Daltonik GmbH, Bremen, Germany). Total number of plasma samples analyzed per group were n = 29 for the LPS group, n = 22 for the IS group and n= 25 for the CTL group. Plasma samples were analyzed with the standard method for microbial biotyping. Briefly, 1μl of plasma was loaded onto each spot in duplicate and 1 μl of the matrix (alpha-cyano-4-hydroxycinnamic acid matrix in 50% acetonitrile and 2.5% trifluoroacetic acid) was added to each dried spot.

Continuous mass spectra were obtained with a Microflex LT/SH MALDI-TOF mass spectrometer using the flexControl software version 3.4.135.0. Acquisition conditions were ionization mode: LD+, acquisition method: MBT_FC.par, acquisition mode: qsim, tof Mode: linear, acquisition Operator Mode: linear, and digitizerType: Bruker BD0G5, within a mass range of 2,000-20,000 Da. Spectra were obtained in the manual mode, using 60% of laser intensity with 40 laser repetition in each shot and reaching between 400-500 spectra by acquisition. Internal calibration was performed every day following manufacturer’s instructions (bacterial test standard; Bruker Daltonik GmbH).

### 2.5 MALDI-TOF-MS data pre-processing

We generated a dataset containing 152 mass spectra from plasma samples corresponding to the three mice groups in duplicate. Mass spectra were read as fid/aqus files with MALDIquantForeign (v0.10) (Gibb, 2019) and they were processed using MALDIquant (v1.16.2) (Gibb and Strimmer, 2012) R package. Briefly, spectra were square root-transformed, smoothed using the Savitzky-Golay algorithm, and were baseline-corrected applying the SNIP process across 100 iterations.

The peaks were detected by a function that estimates the noise of mass spectrometry data by calculating the median absolute deviation (MAD). The signal-to-noise-ratio (SNR) was set up in 4, with a half Window Size of 40 and a tolerance of 0.2 for peak binning. Duplicates were averaged except from one replicate corresponding to the CTL group that was removed due to low quality and 76 averaged spectra were subject to further analysis. Peaks that occurred in less than 33% of the spectra were removed. MALDI-TOF-MS data were transformed by a categorization of the peak intensity, performed with the Binda R package (Gibb, 2015) which compares the intensity value with the peak group average, returning 1 when it was equal or higher and 0 when it was lower. The analysis workflow is depicted in the Figure 1.

**Figure 1.**
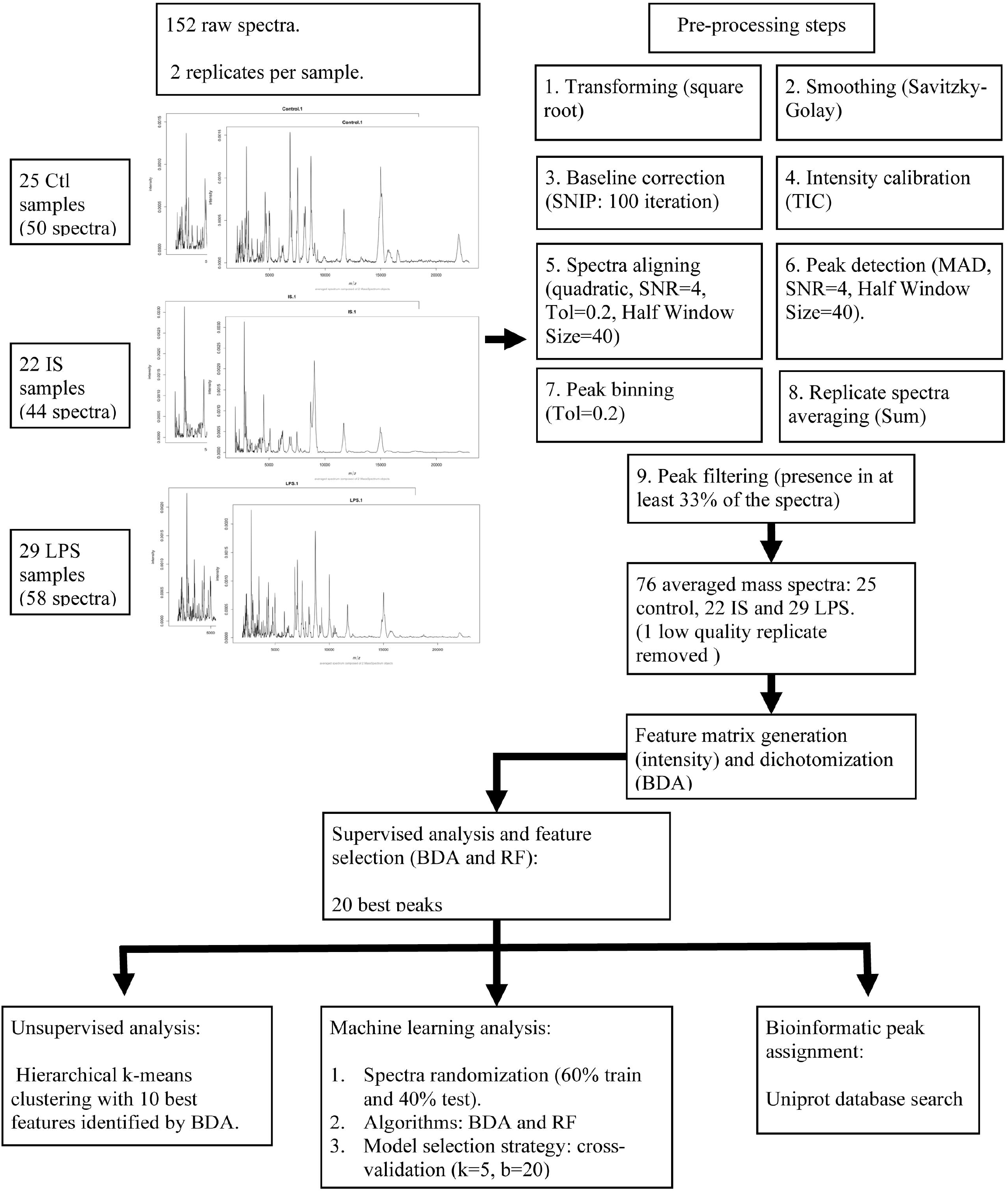
MALDI-TOF- MS data analysis pipeline. Abbreviations. CTL: control group; IS: immunosuppression/anti-inflammatory group; LPS: pro-inflammatory group; BDA: binary discriminant analysis; RF: random forest.

### 2.6 Statistical analysis

#### 2.6.1 Immunological, hematological and biochemical parameters

Graph Pad Prism 6 software (GraphPad Software, San Diego, CA) was used for statistical analysis and plotting. The number of mice or biological replicates (n) analyzed in each experiment were the replicates considered for statistical analysis. Values are expressed as the mean ± standard error of the mean (SEM) of n samples. No outliers were removed. The assumption test to determine the Gaussian distribution was performed by the Kolmogorov and Smirnov method. For parameters with a Gaussian distribution, the differences between two experimental groups were assessed by unpaired Student’s t test, and for multiple group comparisons, as the differences in peripheral blood leukocytes, were analyzed using a one-way ANOVA followed by a Tukey’s multiple comparison test. For parameters with a non-Gaussian distribution, comparisons between two experimental groups were performed using Mann–Whitney U test. All statistical tests were interpreted in a two-tailed fashion and a P<0.05 was considered statistically significant.

#### 2.6.2 MALDI-TOF-MS data

Statistical analysis and plotting were performed using Rstudio. Programmed peaks selection was performed in the entire dataset seeking for biomarkers of each class representing the different experimental groups by the Binary Discriminant Analysis (BDA) algorithm, which outputs the t.score (Class means vs. Pooled mean) of each peak. The sign of the t.score indicates the presence (positive t.score) or absence (negative t.score) of that peak in each group. A significance level of 95% was achieved if the modulus of the t.score was equal or higher than 2.5. The best-extracted features were then used to perform a hierarchical k-means clustering-principal component analysis (PCA) with the factoextra R package (Kassambara and Mundt, 2019). The binary distance was used to measure dissimilarity between observations by the ward.D2 agglomeration method with a k of 4. In addition, spectra were analyzed with the random forest (RF) classifier to test another classification model based on a different underlying criterion.

Afterwards, two ML methods were applied in the dataset to train both BDA and RF classifiers in R (Liaw and Wiener, 2002). Initially, the dataset was randomly partitioned into a training set (60% of plasma samples) and a test set (40% of plasma samples). Programmed feature (peaks) selection was performed with the respective algorithms in the training set seeking for discriminant peaks corresponding to each experimental group. The extracted features were then used to train several ML models. Accuracy, sensitivity, and specificity were used to evaluate the performances of all resulting models in the test set by a cross-validation strategy.

## 3 Results

### 3.1 Simulation of pro- and anti-inflammatory sepsis phases

The proinflammatory phase was induced with a high dose of LPS (LPS group), whereas the anti-inflammatory/immunosuppression phase (IS group) was reached with a scheme of increasing doses of LPS (Suppl. Fig.1). In order to validate the two models, immunological, inflammatory and tissue damage markers were studied in the two experimental groups as well as in the control group (CTL group). In accordance with our previous results, the LPS group was characterized by a marked leukopenia (Suppl. Fig 2a), high plasma levels of pro-inflammatory cytokines such as TNF-α but also of the anti-inflammatory cytokines IL-10 and TGF-β (Suppl. Fig 3a, b). Tissue enzymes indicative of damage were also elevated (Suppl. Fig 4a-c).

On the other hand, the IS group showed a leucocytosis due to increased numbers of circulating granulocytes, particularly of polymorphonuclear neutrophils (Suppl. Fig 2b) (Córdoba-Moreno et al., 2019), along with high plasma levels of TGF- β and low levels of pro-inflammatory cytokines (Suppl. Fig 3c). No signs of tissue damage were observed in this group (Suppl. Fig 4d-f). As previously described (Rearte et al., 2010b; Rearte et al., 2014; Montagna et al., 2020), IS mice had a profound immunological impairment, accompanied by a marked lymphopenia and an increased number of neutrophils in the spleen, which expressed higher levels of the inhibitory receptor PDL-1 (Suppl. Fig 5a, 5b). Moreover, the IS group showed significantly lower antibody titers upon immunization with SRBCs (Suppl. Fig 5c). Collectively, these results indicate that the pro- and anti-inflammatory/immunosuppression phases were emulated in our murine models.

**Figure 2.**
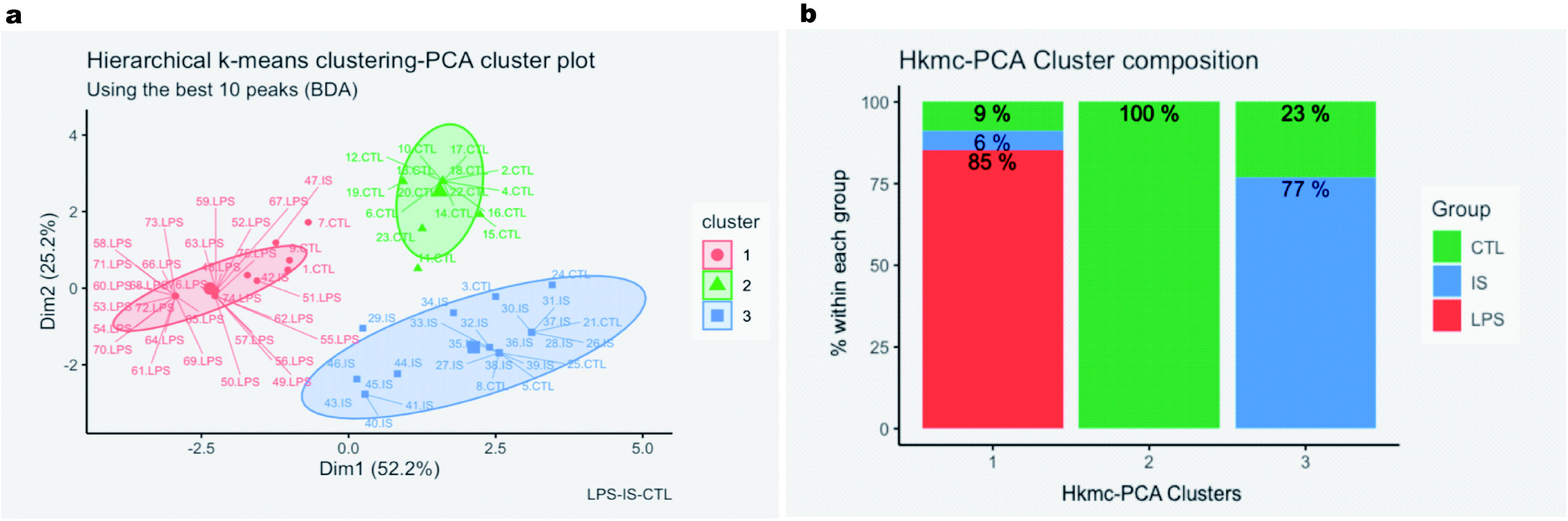
Unsupervised statistical analysis. (**a**) Hierarchical k-means clustering (Hkmc)-Principal Component Analysis (PCA) cluster plot using the top ten peaks selected by the binary discriminant analysis (BDA) algorithm. Labels contain the mice ID and the experimental groups. PC1 (Dim1, x-axis) and 2 (Dim2, y-axis) are depicted. Spectra were clustered into three groups using the Hkmc algorithm, which are represented with three different colors. 95% confidence ellipses were added around cluster means, assuming a multivariate normal distribution. (**b**) Hkmc-PCA cluster composition. The green color represents the CTL mice group, the blue represents the IS group, and the red color represents the LPS group.

### 3.2 MALDI-TOF MS data analysis

Plasma samples of the CTL, IS and LPS groups were analyzed by MALDI-TOF MS in the 2000 – 20,000 Da range. Representative spectra of each experimental group are shown in Suppl. Fig. 6 and the analysis pipeline is detailed in Figure 1. A statistical analysis set was implemented to test if MALDI-TOF MS data were useful to discriminate the three experimental groups. First, through a supervised classification algorithm we looked for peaks that best differentiated the CTL, LPS, and IS groups. The 20 best discriminant peaks identified by binary discriminant analysis (BDA) and random forest (RF) algorithms are shown in the Suppl. Fig. 7a and b respectively, which partially overlapped. Of the 20 best peaks selected by the BDA algorithm, the 10 most discriminant peaks were chosen for an unsupervised statistical analysis to plot the hierarchical k-means clustering-PCA clusters (Figure 2a). Clusters appeared as three non-superimposed groups, and the first two principal components explained 76.6% of the variation with good intra cluster homogeneity. Specifically, cluster 2 achieved 100% (16/16) of homogeneity for spectra corresponding to CTL plasma and 64% (16/25) of CTL spectra fell in this cluster, while for cluster 1 an 85% (29/34) homogeneity was achieved for spectra corresponding to the LPS group and 100% (29/29) of LPS spectra fell in this cluster (Figure 2b). Lastly, homogeneity of cluster 3 was 77% (20/26) mostly represented by spectra of the IS group, with 91% of the IS spectra (20/22) included in this cluster. This result shows that the immunosuppression/anti-inflammatory and pro-inflammatory phases have distinctive plasma peptidome signatures that allow their discrimination.

### 3.3 Machine learning analysis

ML algorithms were used to develop a predictive model based on the plasma fingerprints, using a training data set (14 CTL, 13 IS and 15 LPS samples) challenged with a test data set (11 CTL, 9 IS and 14 LPS samples; Figure 1). To find the most refined model, a dimensionality reduction was performed through the selection of different numbers of distinctive peaks in the training set to train the algorithms, and the performances on the test set were computed for each condition. Specifically, the top 5, 10, 15, and 20 discriminant peaks for BDA and RF algorithms were evaluated by cross-validation (CV; 5 folds and 20 repetitions) and percentages of accuracy, sensitivity, specificity, negative predictive value (NPV) and positive predictive value (PPV) in the test data set were obtained (Table 1).

**Table 1.** Performance of the classification models with the top 5, 10, 15, and 20 peaks.

| Peaks | Algorithm | A (%) | S (%) | E (%) | PPV (%) | NPV (%) |
| --- | --- | --- | --- | --- | --- | --- |
| 5 | BDA | 90.6 | 95.7 | 80 | 90.9 | 89.8 |
| 10 | BDA | 85.6 | 92 | 72.3 | 87.4 | 81.1 |
| 15 | BDA | 87.6 | 93.9 | 74.5 | 88.5 | 85.4 |
| 20 | BDA | 90.1 | 94.3 | 81.4 | 91.4 | 87.3 |
| 5 | RF | 88.2 | 95.7 | 72.7 | 88 | 88 |
| 10 | RF | 91.2 | 91.3 | 90.9 | 95.5 | 83.3 |
| 15 | RF | 88.2 | 91.3 | 81.8 | 91.3 | 81.8 |
| 20 | RF | 94.1 | 95.7 | 90.9 | 95.7 | 90.9 |

A = accuracy, S = sensitivity, E = specificity, PPV = positive predictive value, and NPV = negative predictive value.

Between models, RF using the best 20 peaks showed the most consistent values, with sensitivity and specificity values above 90%. The BDA model trained with the best five peaks showed the best performance in terms of detection of positive cases, with a sensitivity above 95% (Table 1); this strategy constitutes the simplest method in terms of both the number of required peaks and computational demands. These results display the potential of plasma peptidome fingerprints to develop predictive models.

## 4 Discussion

Sepsis constitutes a highly heterogeneous syndrome of complex pathophysiology. The underlying immunological alterations that include pro-inflammatory, anti-inflammatory and immunosuppressive components (Rubio et al., 2019) are reflected in septic plasma protein composition and varies over time according to the disease severity (Hayashi et al., 2019).

In the last decades, several circulating markers with recognized functions in sepsis were identified. However, none of these markers showed the specificity and sensitivity required for its clinical use (Parlato et al., 2018; van Engelen et al., 2018b). Furthermore, methods that rely on immunolabeling such as ELISA or flow cytometry, are highly specific and sensitive for the detection of low abundance markers, but requires costly equipment and consumables, as well as specific technical knowhow. In addition, these techniques are time consuming, limiting their clinical usefulness for early decision-making.

Regarding biomarker discovery, proteomics proved to be an important source of information. However, in the classical approaches the plasma samples are subjected to depletion of abundant proteins (Ludwig and Hummon, 2017) and fractionation to identify individual biomarkers (Hayashi et al., 2019; Harberts et al., 2020) which requires more analysis time, specific equipment and higher costs. These constraints led us to explore a top-down strategy that could overcome these difficulties. Thus, having in mind that we aimed to design a readily transferrable method, our approach was meant to be as simple as possible and unlabeled whole plasma were directly read by MALDI-TOF-MS.

Our results showed that the performance of the test with whole plasma and standard acquisition settings used in routine microbiological biotyping were excellent in terms of discriminatory power. First, an ensemble between supervised and unsupervised algorithms allowed the overall discrimination of plasma samples corresponding to the three experimental groups. These results showed that each phase comprise a distinctive MALDI-TOF-MS pattern, which turns this approach into a promising tool to determine different phases in sepsis. A recent study reported a top-down plasma analysis showing that differences between samples from healthy donors and cancer patients could be appreciated without the depletion of large abundant proteins by setting a cut-off at 30KDa (Cheon et al., 2016), supporting our results.

Furthermore, the performance of our ML algorithms challenged with the test dataset, was able to identify basal, pro-inflammatory and anti-inflammatory/immunosuppressive phases in a very satisfactory way, exceeding 90% of accuracy in some of the tested conditions. The cross validation strategy used herein allows a fine-tuning of the models according to particular needs. Thus, depending on the clinical and epidemiological settings, algorithms with higher sensitivity and lower computational demand like our 5-peaks BDA model could be preferred over those with a good overall accuracy.

The peaks found in the 2-20KDa range mostly correspond to small peptides. Plasma peptidome has been proposed as an interesting source of information for diagnostic purposes (Cheon et al., 2016; Dufresne et al., 2018). These peptides are presumably fragments of larger proteins derived from digestion by proteases or non-enzymatic degradation (Greening and Simpson, 2017) but the lack of comprehensive digestome/degradome databases is still a limitation to their study (Shen et al., 2010). Nevertheless, recent reports demonstrate the importance of plasma peptidome as a useful tool for clinical monitoring in septic shock (Aletti et al., 2016; Bauza-Martinez et al., 2018). Although the identification of individual peaks is beyond the objectives of this study, it would be interesting to evaluate them in the future with specific methods with higher resolving power.

A major limitation of this study is related to the animal models chosen. Although LPS-induced inflammation/immunosuppression models are widely used, they do not accurately represent the dynamic physiological changes that occur in sepsis (Lewis et al., 2016). Endotoxin challenge promotes a faster and transient release of pro/anti-inflammatory mediators compared to the septic processes. However, these models allowed us to define two discrete phases that actually occur in a septic process. For this reason, we considered this was the most appropriate approach to test the potential of MALDI-TOF-MS analysis in the proof-of concept phase. We are currently validating our results in a dynamic model of sepsis, the cecal ligation and puncture model, with very encouraging results.

Our approach has numerous advantages. MALDI-TOF-MS is a simple and speedy method, a critical feature considering the dynamism of sepsis, which is spreading in the clinical setting even in resource-constrained regions. In addition, we used an accessible sample as plasma, without complex pre-processing steps. Finally, we performed our ML analysis in the open source software R that also guarantees the transferability of the bioinformatics tool.

In conclusion, our results indicate that plasma peptidome analysis by MALDI-TOF-MS has the potential to be a highly relevant strategy for sepsis patient stratification that could constitute a powerful tool for therapeutic decision making in sepsis depending on the pro- or anti-inflammatory phases the patient is undergoing.

## 5 Conflict of Interest

The authors declare that the research was conducted in the absence of any commercial or financial relationships that could be construed as a potential conflict of interest.

## 6 Author Contributions

ML designed and performed the bioinformatic analysis. BR and NY conceived and designed the study. MFT, LM and BR performed the experiments with mice and analyzed data. MP and CV provided materials and technical support. MG and BL provided materials and collaborated in study design. ML and NY performed data curation. NY and BR analyzed and discussed the results. BR was responsible for funding acquisition and study supervision. Manuscript was written by NY and BR, and was corrected by ML and BL. All the authors revised and approved the final version of the manuscript.

## 7 Funding

This study was supported by the Agencia Nacional de Promoción Científica y Tecnológica [grant number PICT 2015-0412], Buenos Aires, Argentina.

## 8 Acknowledgments

We thank Dr. Martín Isturiz for his eternal accompaniment and support for the development of this study. His outstanding contribution stems from his singular thoughts. Forever in our hearts, Dr. Isturiz. We also thank Dr. Viviana Ritacco for her thoughtful comments on the manuscript. A preliminary version of the manuscript was deposited in the preprint server BioRXiv (doi: 10.1101/2020.12.29.424724).

## 9 Data Availability Statement

The MS datasets generated for this study as well as the analysis pipelines can be found in GitHub [https://github.com/MarManLed/SepsisData].

## Notes

### Competing Interest Statement

The authors have declared no competing interest.

