## Supplementary Figures for "Peptidome profiling for the immunological stratification in sepsis: a proof of concept study"

Supplementary Material

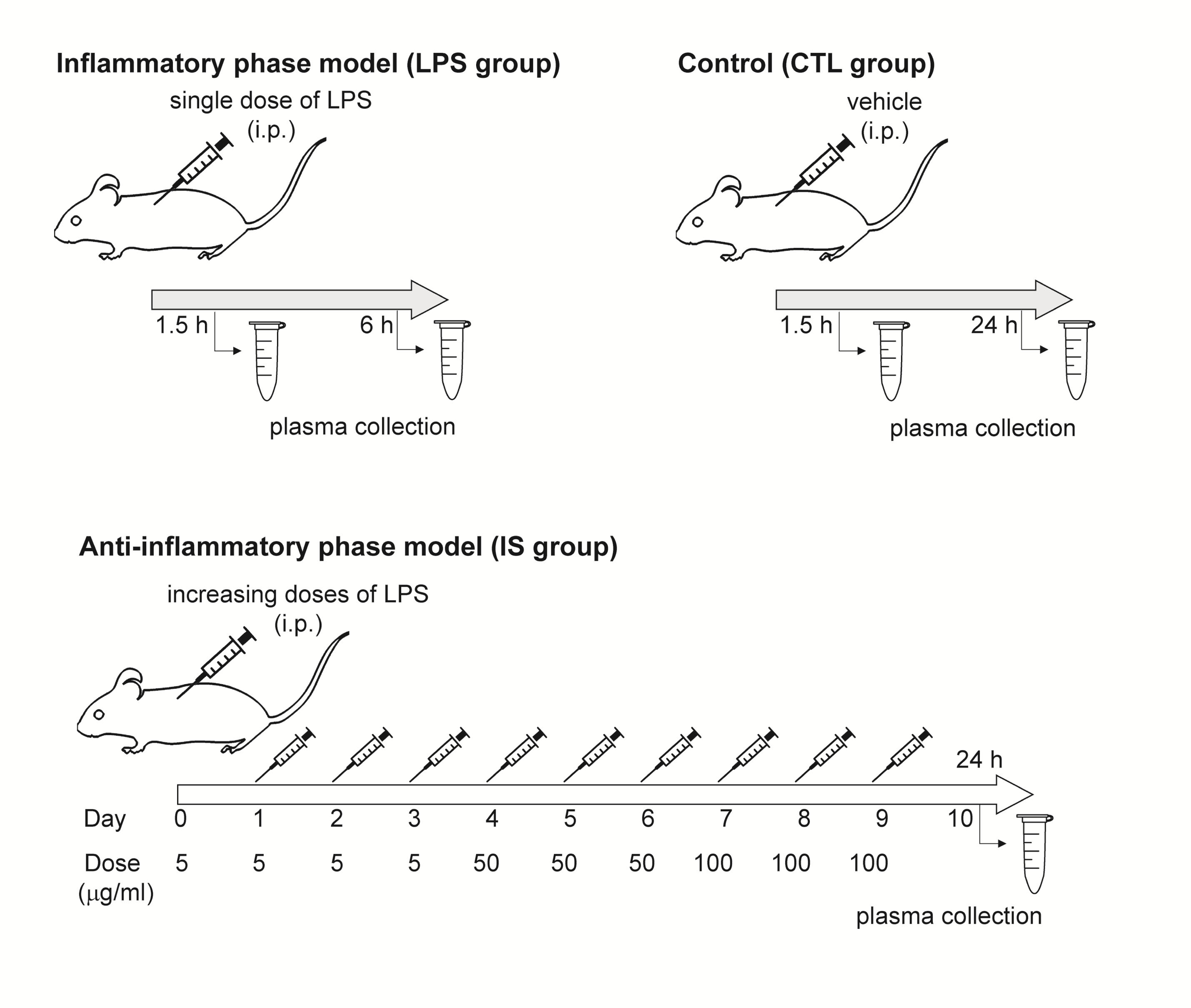

**Supplementary figure 1.** Inoculation and sample collection schemes. Inoculation schemes of the proinflammatory (LPS) group, the anti-inflammatory/immunosuppression (IS) group and the basal control (CTL) group. Plasma were collected in heparinized tubes at the indicated time points through submandibular bleeding. i.p.: intraperitoneally.

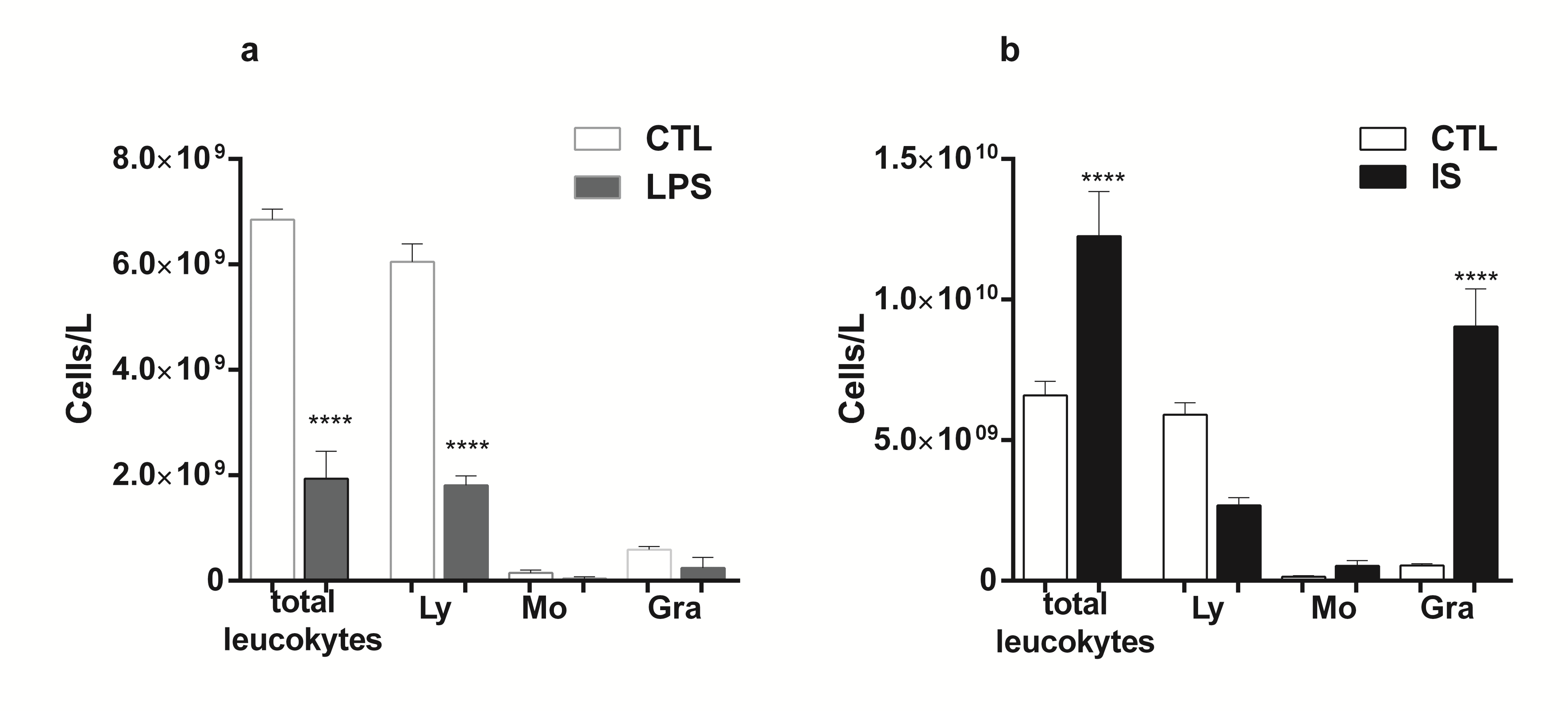

**Supplementary figure 2.** Peripheral blood leukocytes count. BALB/c mice were inoculated with one LPS dose (pro-inflammatory phase; LPS group) (**a**), or with successive and increasing LPS doses (immunosuppression phase; IS group) (**b**). Inoculation and blood collection schemes are detailed in the Suppl. Fig. 1. Blood was collected after 1.5 h (LPS group) or 24h after the last LPS dose LPS (IS group). A control group (CTL) was inoculated with vehicle (saline solution) and the plasma was collected at the same time points. The samples were analyzed with a Coulter hematology analyzer. Results are expressed as the mean ± SEM; n= 6 to 7 per group. Data are representative of two independent experiments. ****P<0.0001 compared to the same cell type in the CTL group. One-way ANOVA and Tukey’s multiple comparisons test. Ly: lymphocytes; Mo: monocytes; Gra: granulocytes.

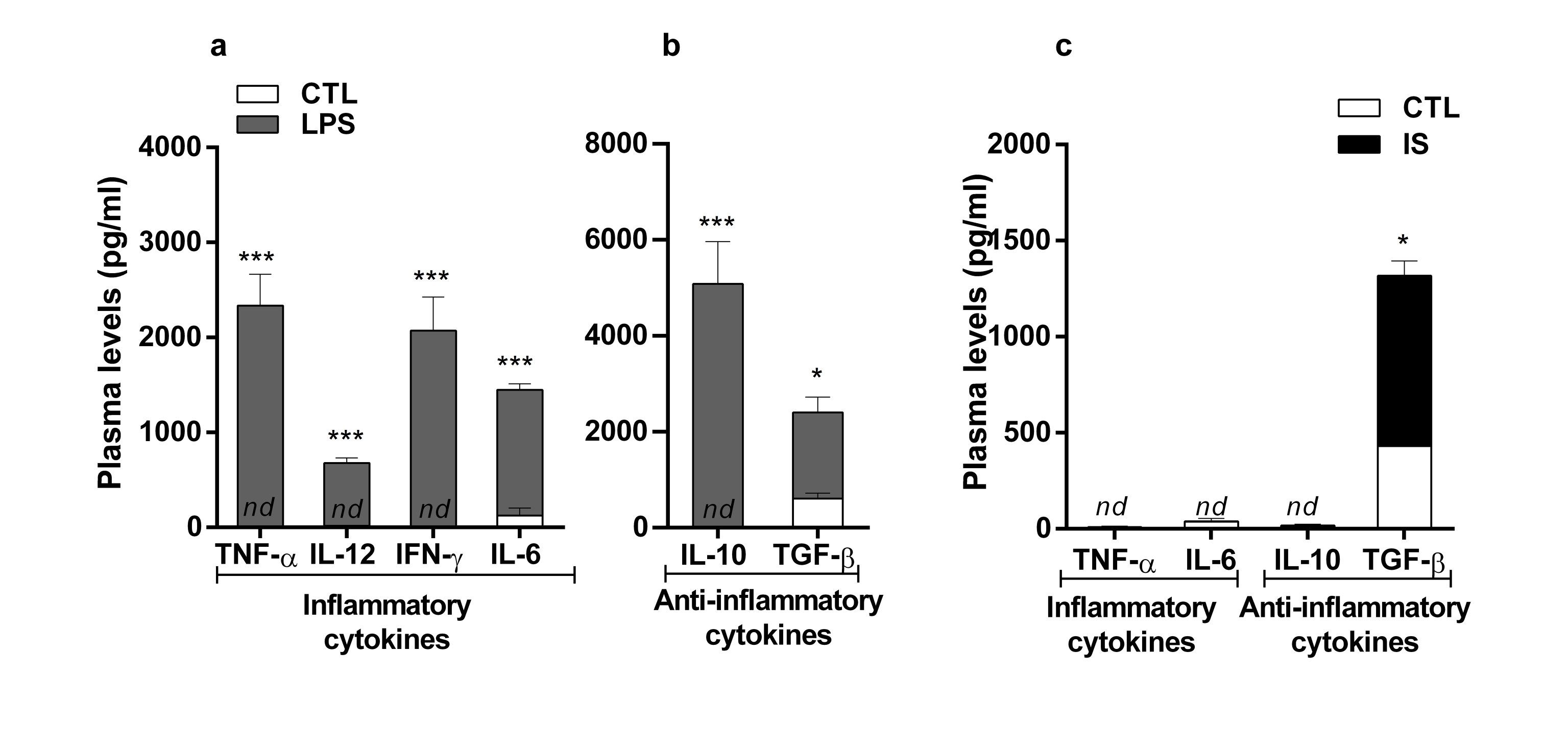

**Supplementary figure 3.** Pro and anti-inflammatory cytokines levels in plasma. BALB/c mice were inoculated with one LPS dose (pro-inflammatory phase; LPS group) (**a, b**), or with successive and increasing LPS doses (immunosuppression phase; IS group) (**c**). At different times after the last LPS challenge plasma were collected, and the cytokines levels were evaluated through an ELISA. (**a, b**) TNF-α, IL-6, IL-10 and TGF-β were evaluated 1.5h post LPS; IL-12 and IFN-γ at the 6h post LPS challenge. (**c**) Cytokines TNF-α, IL-6, IL-10 and TGF-β were evaluated 24h after the last LPS dose. A control group (CTL) was inoculated with vehicle (saline solution) and the plasma was collected at the same time points. Results are expressed as the mean ± SEM; n= 6 to 7 per group. Data are representative of two independent experiments. *P<0.05, **P<0.01, ***P<0.001, ****P<0.0001 compared with CTL mice at the same time; Mann-Whitney test.

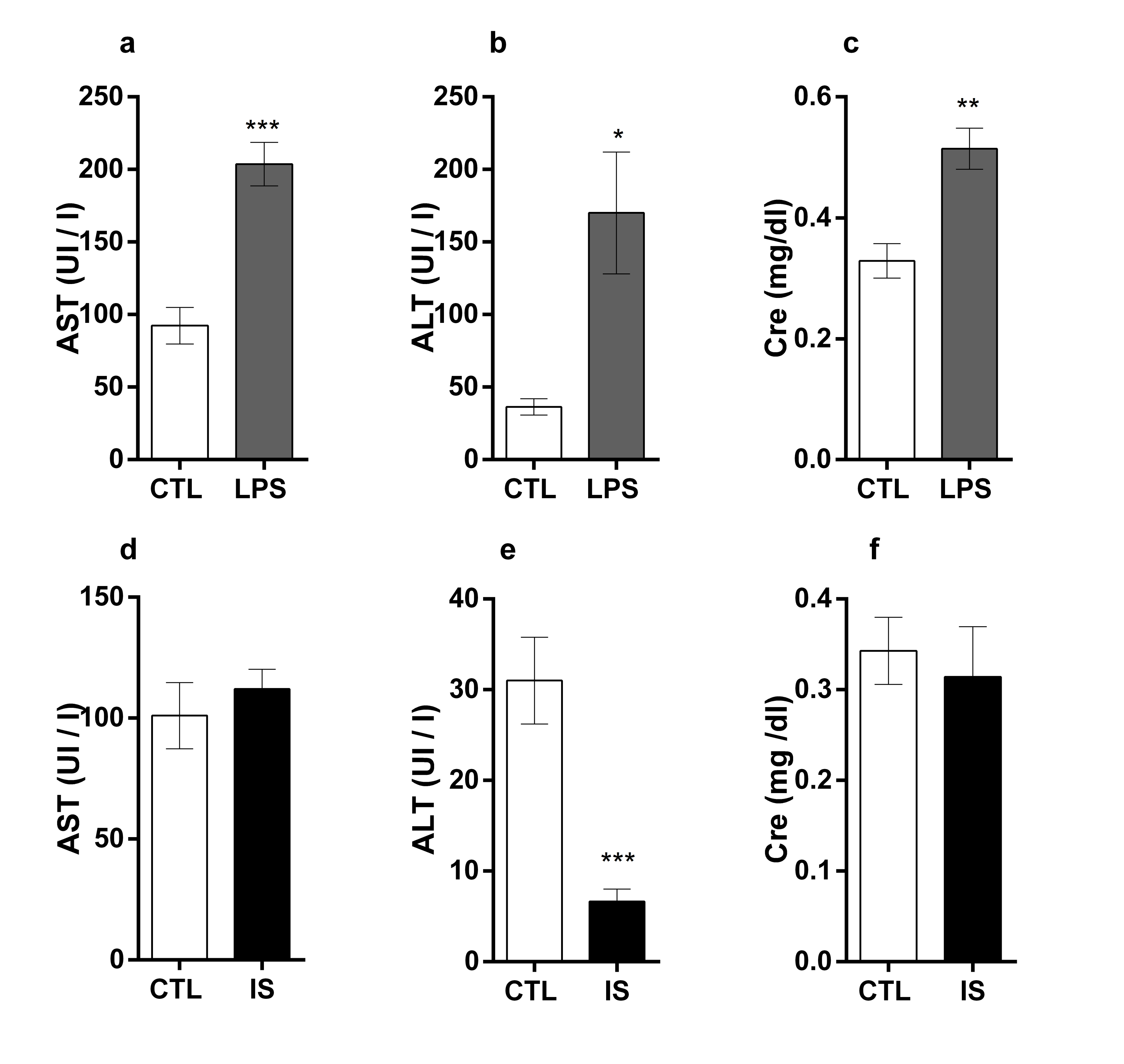

**Supplementary figure 4.** Tissue damage indicators in plasma. BALB/c mice were inoculated with one LPS dose (pro-inflammatory phase; LPS group) (**a, b, c**), or with successive and increasing LPS doses (anti-inflammatory/immunosuppression phase; IS group) (**d, e, f**). After 6 h (LPS group) or 24 h (IS group) after the last LPS administration, plasma were collected and enzyme levels were evaluated. A control group (CTL) was inoculated with vehicle (saline solution) and the plasma were collected at the same time points. AST: aspartate transaminase; ALT: alanine transaminase; Cre: creatinine. Results are expressed as the mean ± SEM; n= 5 to 7 per group. Data are representative of two independent experiments. *P<0.05, **P<0.01, ***P<0.001, ****P<0.0001 compared with CTL mice at the same time; Student’s t-test.

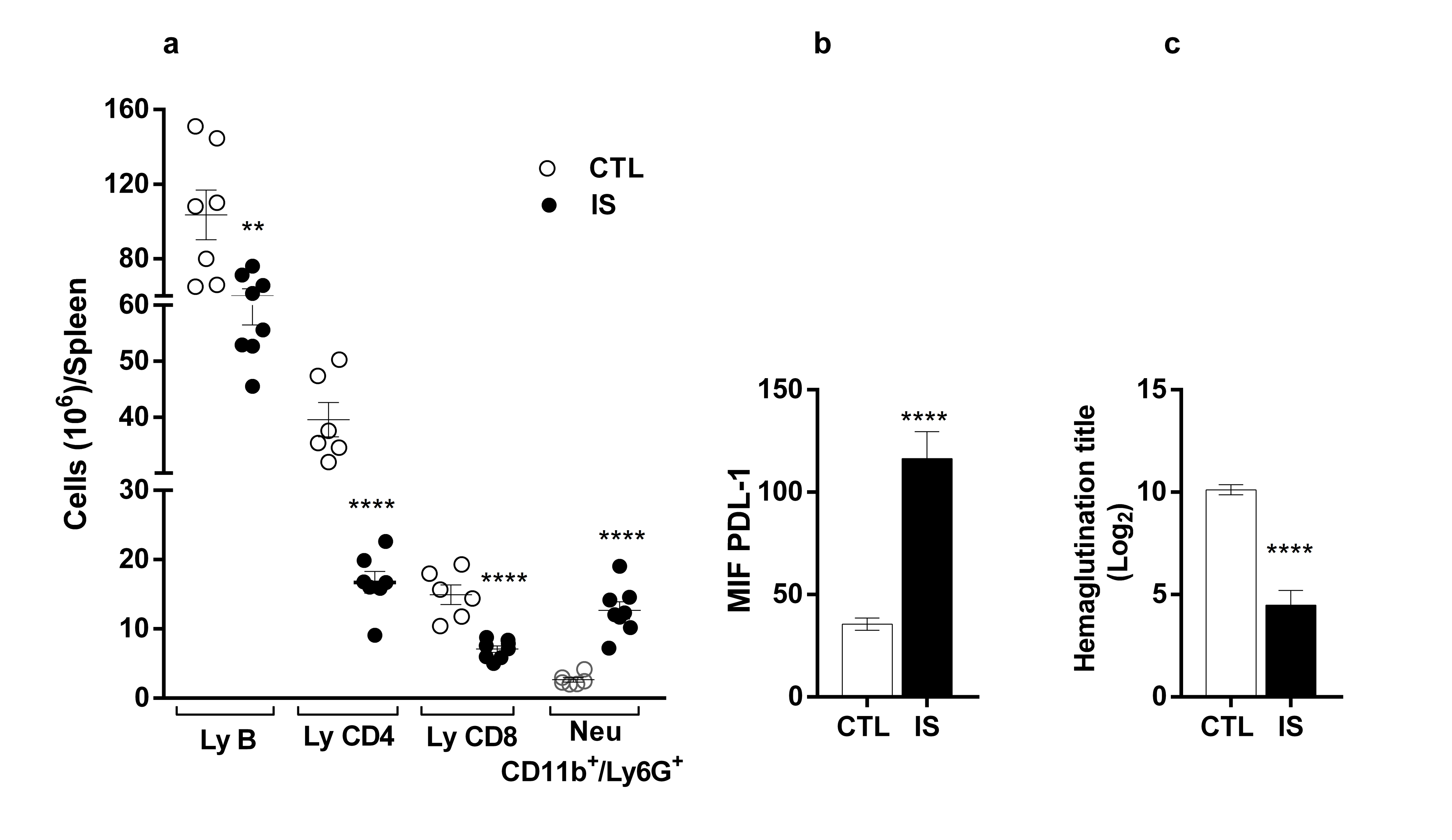

**Supplementary figure 5.** Lymphocyte and myeloid cell populations in spleen and humoral immune response during the immunosuppression phase. BALB/c mice were inoculated with successive and increasing LPS doses (immunosuppression phase; IS group). A control group (CTL) was inoculated with vehicle (saline solution). 24h after the last LPS dose or vehicle, the spleen were removed and the cellular suspension was evaluated by flow cytometry. (**a**) Total numbers of B lymphocytes (B Ly), CD4 and CD8 T lymphocytes (Ly CD4; Ly CD8) and neutrophils (Neu; CD11b/Ly6G) from the IS and CTL spleen were evaluated. Ly and myeloid gates were defined by features of forward and side scatter. (**b**) PDL-1 expression on splenic CD11b myeloid cells of IS and CTL mice was evaluated. Results are expressed as the mean ± SEM; n= 6 to 8 per group. Data are representative of two independent experiments. **P<0.01, ****P<0.0001 compared with to the same cell type in the CTL group; Student’s t-test. (**c**) BALB/c mice from IS were immunized with sheep red blood cells (SRBCs; 5x10^8^/mouse, 0.1 ml i.p.) 24 h after the last LPS dose. CTL mice were immunized with the same antigen. Seven days after the immunization, the mice were bled and the serum was collected. The anti-SRBC antibody titer was evaluated by hemagglutination assay. Results are expressed as the mean ± SEM; n= 8 per group. Data are representative of two independent experiments. ****P<0.0001 compared with the CTL group; Student’s t-test.

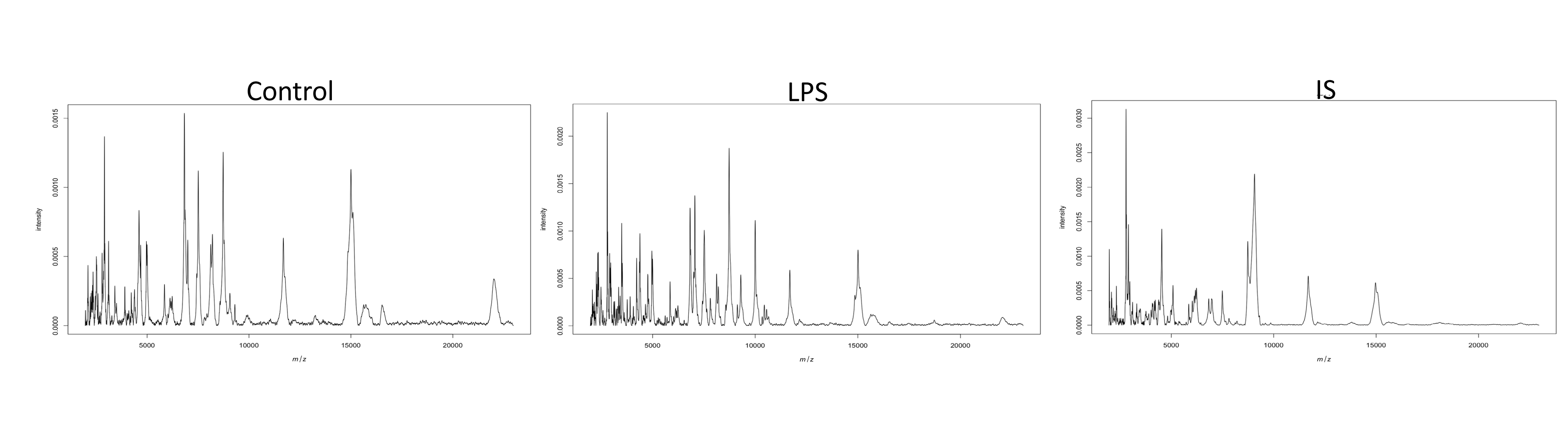

**Supplementary figure 6.** Representative average mass spectra of each experimental group. The y-axis represents the calibrated intensity of each detected peaks, which are represented in the x-axis. Each image represents the average of 2 mass spectra acquired in duplicate.

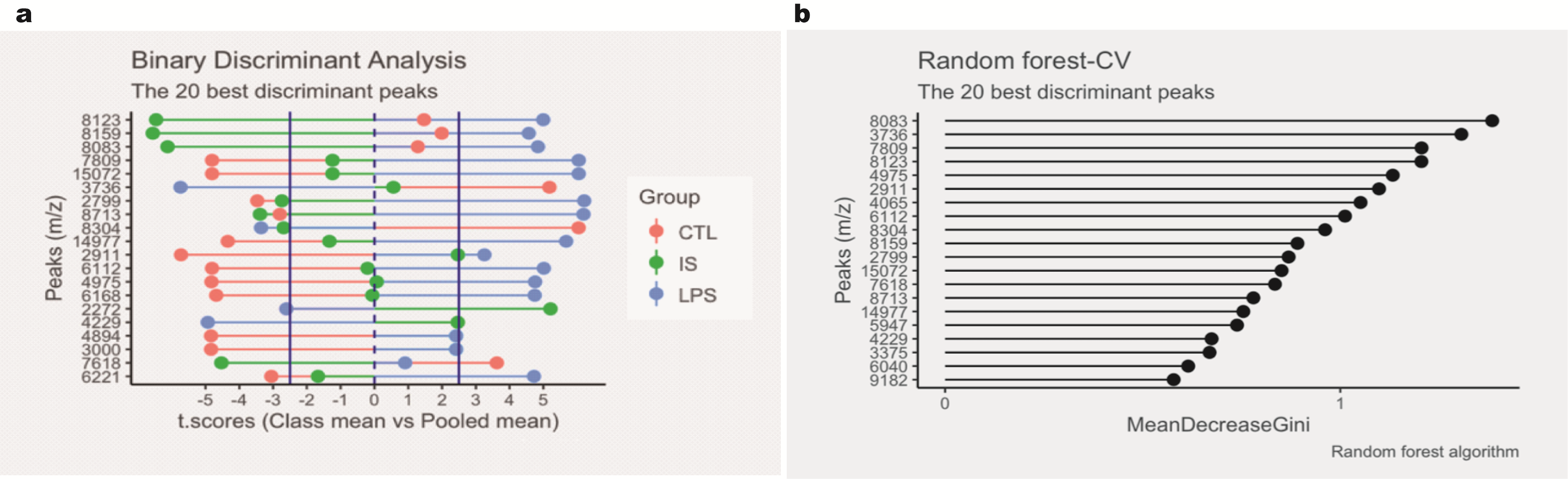

**Supplementary figure 7.** Feature selection plots. The 20 best peaks identified by each algorithm were selected. (**a**) Binary discriminant analysis (BDA) peak selection. The algorithm outputs the t.score (x-axis = Class means vs. Pooled mean) of each peak (y-axis). The sign of the t.score indicates the presence (positive t.score) or absence (negative t.score) of that peak in each group. A significance level of 95% was achieved if the t.score was equal or higher than 2.5 and equal or less than -2.5. (**b**) Random forest (RF) peak selection. The algorithm outputs the mean decrease in the Gini index, which is plotted in the x-axis, by each feature represented in the y axis.
